# Selective behavioural impairments in mice heterozygous for the cross disorder psychiatric risk gene *DLG2*

**DOI:** 10.1101/2021.10.05.463181

**Authors:** Rachel Pass, Niels Haan, Trevor Humby, Lawrence S. Wilkinson, Jeremy Hall, Kerrie L. Thomas

## Abstract

Mutations affecting *DLG2* are emerging as a genetic risk factor associated with neurodevelopmental psychiatric disorders including schizophrenia, autism spectrum disorder and bipolar disorder. Discs large homolog 2 (DLG2) is a member of the membrane-associated guanylate kinase protein superfamily of scaffold proteins, a component of the post-synaptic density in excitatory neurons and regulator of synaptic function and plasticity. It remains an important question whether and how haploinsuffiency of DLG2 contributes to impairments in basic behavioural and cognitive functions that may underlie symptomatic domains in patients that cross diagnostic boundaries. Using a heterozygous *Dlg2* mouse model we examined the impact of reduced *Dlg2* expression on functions commonly impaired in neurodevelopmental psychiatric disorders including motor co-ordination and learning, pre-pulse inhibition and habituation to novel stimuli. The heterozygous *Dlg2* mice exhibited behavioural impairments in long-term motor learning and long-term habituation to a novel context, but not motor co-ordination, initial responses to a novel context, PPI of acoustic startle or anxiety. We additionally showed evidence for the reduced regulation of the synaptic plasticity-associated protein cFos in the motor cortex during motor learning. The sensitivity of selective behavioural and cognitive functions, particularly those dependent on synaptic plasticity, to reduced expression of DLG2 give further credence for DLG2 playing a critical role in specific brain functions but also a mechanistic understanding of symptom expression shared across psychiatric disorders.

## Introduction

Schizophrenia (SZ), autism spectrum disorder (ASD) and bipolar disorder (BD) are often comorbid neurodevelopmental psychiatric disorders ^1,2^, with overlapping symptoms, including cognitive, neurobehavioural and motor dysfunction ^3–5^. There is also a substantial portion of shared genetic risk^6–8^, concordant with the correlation between functional disabilities and risk genes associated with developmental disorders ^9,10^. Genetic analysis approaches, including genome wide association studies (GWAS), studies of common variants as well as copy number variant (CNV) analysis, and exome sequencing for rarer but more penetrant mutations, combined with subsequent gene set enrichment analysis have been important for identifying the biological pathways underlying disorder pathophysiology and common targets for clinical interventions ^11^. This approach has uncovered an enrichment of genes involved in synaptic plasticity and the post-synaptic density (PSD) in SZ ^6,12–17^. Convergence of mutations in genes important for synaptic formation, plasticity and elimination have also been found in ASD ^18–22^,with meta-analyses identifying postsynaptic complexes as one of three functionally interconnected pathways associated with ASD risk genes ^23,24^.

Discs large homolog 2 (DLG2, also known as post-synaptic density protein-93 (PSD-93) and chapsyn-110), is part of the membrane associated guanylate kinase (MAGUK) protein superfamily of scaffold proteins and is enriched at the PSD ^25^. MAGUKs primarily bind and stabilise proteins at synapses, and regulate the synaptic localization of glutamate receptors, key for synaptic transmission and synaptic plasticity ^26^. Both *de novo* CNVs and SNVs implicate *DLG2* in increasing risk of schizophrenia ^6,15,27–29^, ASD ^30–32^, and potentially BD ^33^.

Mutations in *DLG2* affecting its function and/or expression are plausibly related to the impairments of learning and memory underlying in the cognitive deficits seen in SZ, as well as altered synaptic plasticity linked to abnormal sociability and motor deficits reminiscent of ASD ^22,34–37^. Indeed, homozygous knockout mice (*Dlg2*^−/−^) showed no impairment in simple forms of learning, but showed deficits in complex learning, cognitive flexibility and attention comparable to similar deficits in humans carrying mutations disrupting the coding region of *DLG2* ^38^. Furthermore, *Dlg2*^−/−^ mice displayed abnormal social behaviours ^39,40^, increased repetitive behaviours and hypoactivity in response to novelty ^40^, as well as defective long-term potentiation (LTP) ^41^ and aberrant excitatory synaptic transmission in the dorsal striatum ^40^. Significantly, the functions of Dlg2 and other members of the Dlg family are dissociable both at the behavioural and synaptic levels ^26,41–43^.

Most genetic lesions observed in humans are deletions within or of *DLG2* ^15,27–29,44^ and are heterozygous ^32,45^. Little attention has been paid to investigating behavioural phenotypes associated with heterozygous *Dlg2* models, arguably more translationally relevant than homozygous knockouts. Winkler and colleagues (2018) identified a *Dlg2* dose-dependent impairment of motor learning in mice. There has been no systematic investigation of reduced *Dlg2* gene dosage on potentially core endophenotypes associated with SZ, ASD, BD and other neuropsychiatric disorders including motor function, pre-pulse inhibition (PPI) of startle responses and habituation to a novel stimulus ^46–55^.

In this study we investigated motor co-ordination and motor learning, habituation to a novel context, PPI of the acoustic startle response and anxiety in heterozygous *Dlg2* adult male mice. Using cFos levels as a proxy marker for synaptic activity and plasticity ^56^, we also tested the hypothesis that in the heterozygous *Dlg2* mice impaired behavioural function would be correlated with reduced activity-dependent cFos activation.

## Methods

### Animals

All procedures were conducted in accordance with local ethical guidelines and Animals (Scientific Procedures) Act (ASPA) (1986) under UK Home Office project license PPL 30/3135. *Dlg2*^tm1a(EUCOMM)Wtsi^ mice (MGI:4842622, hereafter *Dlg2*^+/−^ or HET) were maintained heterozygously on a C57/BL6J background. Genotyping was conducted in house (*Supplementary Methods*). *Dlg2*^+/−^ and WT controls were housed in mixed-genotype standard cages with ≤ 5 littermates. Mice were maintained on a 12:12 hour light/dark cycle (light phase 8am – 8pm) with *ad libtum* access to standard food (RM3 E, Special Services Diet, Lillico, UK) and water. Cages were lined with wood shavings, and cardboard tubes and wooden sticks provided environmental enrichment. Holding rooms were maintained at 45-60% humidity and 19-22°C.

### Behavioural analysis

Data from three separate male cohorts is presented in this manuscript (Supplementary Table 1). At start of testing mice were 8-12 weeks of age. All tasks were completed prior to lights off. All mice were habituated to the experimenter for one week before testing, and to procedural rooms for 5 minutes before training and testing.

### Motor learning and performance

#### Two-day (2d) rotarod motor learning and motor performance task

Mice were habituated to the experimental room for 5 minutes prior to the first trial. Each mouse received 5-minute trials on the rotarod (3 cm drum separated by flanges into 5 × 5.7 cm individual lanes, fall height 16 cm, 47600, Ugo Basile, Italy). Motor training comprised of 5 trials across two days (Day 1, two trials, Day 2, three trials) during which the rotarod was accelerated incrementally from 5-50 rpm over 5 minutes. Animals were returned to home cages in the testing room between trials. Following completion of a trial for the whole cohort (approx. ITI 30 mins) the next trial was started. Starting on Day, 3 to assess motor performance, mice then completed two 5-minute trials at a fixed speed (5-45 rpm in 5 rpm increments) over 5 days, with an inter-trial interval of 20-30 minutes during which the mice were returned to their home cage in the test room. The mean latency to fall for two trials per speed was calculated.

#### Three day (3d) rotarod motor learning task

Mice were habituated to the experimental room for 5 minutes prior to the first trial. Each mouse received 7 trials a day for 3 consecutive days. Each trial comprised of the rotarod accelerating incrementally from 4-40 rpm over 5 minutes, with an inter-trial interval of 5 minutes during which the mice were returned to their home cage in the testing room. For the assessment of cortical activity using cFos IHC, experimental mice were trained on the accelerating rotarod under the same conditions. However, mice were single caged in their housing (not test) room for 1.5 hours after the second trial on Day 1. Control mice were singled caged in the holding room for 1.5 hours.

In all tasks, mice were placed on the rod facing away from the experimenter. Rotarod training order was counterbalanced for genotype. Latency to fall (s), defined as the first fall from the rod or 1 full rotation on the rod, was measured for each mouse per trial. Consolidation of motor learning was analysed by comparing latency to fall between for the last trial on Day 1 (D1T7) to the first trial on Day 2 (D2T1), = latency to fall(s) (D2T1/(D2T1+D1T7).

### Acoustic startle and pre-pulse inhibition (PPI)

Acoustic startle response (ASR) and pre-pulse inhibition (PPI) of startle response was monitored using a SR-Lab™ Startle Response System (San Diego Instruments, USA). Mice were placed in a Perspex tube (internal diameter 35mm) mounted to a Perspex plinth containing a piezoelectric pressure-sensitive accelerometer sensor. All data are reported as the weight adjusted average response amplitude (v avg) and the contribution of body weight to the startle response was accounted for using Kleiber’s 0.75 mass exponent ^57^. A session began with a 5-minute presentation of scrambled white noise at background intensity (70 db) to habituate the animals to the apparatus followed by 3 blocks of acoustic stimuli (pulses). The startle amplitude was 120dB for block 1, 105dB for block 2 and a range (80-120dB) in block 3. Pulse alone trials consisted of a 40ms startle stimulus whilst pre-pulse trials consisted of a 20ms pre-pulse at 4, 8 or 16dB above background followed by a 40ms startle stimulus (120 or 105dB) 70ms after the pre-pulse. Blocks 1 and 2 began with 6 pulse alone trials (either 120dB or 105dB) followed by pseudorandom stimuli (pulse alone, pre-pulse or no stimulus) every 15s. In block 3 varying pulse alone stimuli (80-120dB) were presented 3 times in a pseudorandom manner. Whole body startle responses were recorded as average startle during the 15s window after pulse onset. No stimulus trials separated the blocks and ended the session.

The startle response (startle amplitude) to the first three pulses at 105 dB and 120 dB were averaged and analysed as an index of emotional reactivity before appreciable habituation ^58^. To measure habituation of the startle response, responses to the first five pulse-alone trials at each intensity were measured. For each animal the absolute PPI unconfounded by the initial startle response was calculated as the percentage reduction in startle amplitude in pre-pulse (PPI) compared to pulse alone trials (105 dB and 120 dB). [%PPI=100 × (ASRstartle pulse alone − ASRprepulse + startle pulse)/ASRstartle pulse alone].

### Locomotor activity and habituation

The locomotor activity of individual mice was conducted in clear Perspex activity boxes (CeNeS Cognition, Cambridge, UK, 21cm × 36cm × 20cm; 2 IR beams 3 cm from box end, 1 cm from box floor) in the dark. For each mouse, individual beam breaks were recorded (using a custom BBC BASIC V6 programme with additional ARACHNID interfacing, Campden Instruments, Loughborough, UK) in 5 minute bins over a 2 hr session for 5 consecutive days. Mice were assigned an activity box for the duration of experiment. Twelve mice were run concurrently and the order/time of testing was conserved across days. Each batch of 12 mice was counterbalanced for genotype. The total number of beam breaks per day was recorded and the total number of beam breaks in the first 6 bins (30 min quartile, Q1-Q4) each day was calculated to measure between-session and within-session activity, respectively. Changes in between-session activity = beam breaks (Day 5/(Day1 + Day 5)).

### RNA extraction and RT-qPCR

Mice (12 WT, 12 HET) were culled via cervical dislocation and the pre-frontal cortex (PFC), hippocampus and cerebellum dissected and flash frozen and samples stored at −80 °C until use. RNA from tissue samples was isolated using QIAGEN RNeasy kits and DNase treated with an Ambion TURBO DNA-free™ Kit (ThermoFisher Scientific, Delaware, USA). cDNA synthesis from 1.5ng/μl RNA was performed using RNA to cDNA EcoDry Premix (Random Hexamers) (TaKaRa Europe SAS, Saint-Germain-en-Laye, France). cDNA was diluted to achieve a 1:15 dilution and then stored at −20°C prior to qPCR. qPCR was conducted with the SensiMix SYBR® No-ROX Kit (Meridian Bioscience, Cincinnati USA) on an StepOne Plus Applied Biosystems Real-Time PCR instrument (ThermoFisher Scientific, Delaware, USA;

1 cycle 95°C, 10 mins; 45 cycles of 95°C, 15s; 60°C, 1 min). Potential compensatory changes in the expression of other *Dlgs* genes have been found in homozygous *Dlg2* mice previously ^59^, so mRNA expression of *Dlg1, Dlg3* and *Dlg4* was measured in addition to *Dlg2*. *Hprt* and *Gapdh* acted as housekeeping genes. Primers were designed using NCBI Primer-BLAST (5’-3’: *Dlg1* - CGAAGAACAGTCTGGGCCTT (forward), GGGGATCTGTGTCAGTGTGG (reverse); *Dlg2* - TGCCTGGCTGGAGTTTACAG (forward), TTTTACAATGGGGCCTCCGC (reverse); *Dlg3* - GAGCCAGTGACACGACAAGA (forward), GCGGGAACTCAGAGATGAGG (reverse); *Dlg4* - GGGCCTAAAGGACTTGGCTT (forward), TGACATCCTCTAGCCCCACA (reverse); *Gapdh* - GAACATCATCCCTGCATCCA (forward), CCAGTGAGCTTCCCGTTCA (reverse); *UBC* - CCAGTGTTACCACCAAGAAGGT (forward), CCATCACACCCAAGAACAAGC (reverse) and commercially synthesised (Merck Life Science UK Ltd, Gillingham, Dorset, UK). All qPCR samples were run in triplicate and the outcome was calculated using the 2^−ΔΔCt^ method ^60^.

### cFos immunohistochemistry

Mice were transcardially perfused under terminal anaesthesia (0.1 ml 200mg/ml Euthatal, Duggan Veterinary Supplies Ltd, Tipperary, Ireland) using PBS and 4% PFA and the brains removed. Following 24 hrs post-fixation in 4% PFA, the brains were cryopreserved in 30% sucrose in PBS at 4°C then embedded in Tissue Plus O.C.T. compound (ThermoFisher Scientific, UK) before storage at −80°C. 1 in 10 coronal sections (40 μm) were cut and collected using a Lecia CM1900 cyrostat (Leica Biosystems, UK) spanning the emergence of cortical M1 (approx. Bregma +2.3 to +2.2 mm) and stored in 1X PBS at 4°C.

Sections were blocked in 1% Triton-X, 500 μl PBS (PBST) containing 3% normal donkey serum (S30-100ML, Millipore, Hertfordshire, UK) at room temperature (RT) with agitation for 2 hours. Rabbit anti-cFos (ABE457, 1:5000, Merck Millipore, Hertfordshire, UK) was diluted in 500 μl 0.1% PBST with 0.2% normal donkey serum (v/v) and incubated overnight with agitation at 4°C. Following washes, Invitrogen Alexa Fluor Plus 488® goat anti-rabbit secondary antibody (A32731 ThermoFisher Scientific, UK) was diluted (1:1000) in 500 μl 0.1% PBST with 0.2% normal donkey serum (S30-100ML, Merck Millipore, Hertfordshire, UK) and sections were protected from light and incubated at RT with agitation for 2 hours. Sections were incubated with the nucleus DNA stain *4′,6-diamidino-2-phenylindole* (DAPI) (1μg/ml D9542-10MG, Merck Life Science UK Ltd, Gillingham, Dorset, UK) in 1x PBS at RT with agitation for 5 minutes, then washed. Sections were mounted in a counterbalanced manner, coverslipped with 20 μl Mowiol® 4-88 (Merck Life Science UK Ltd, Gillingham, Dorset, UK) and stored at 4°C.

### Imaging

M1 was imaged as a Z stack using a Zeiss LSM 900 confocal microscope running ZEN 2 (blue) software at 20X magnification and analysis was conducted in Fiji (1.52g, https://imagej.nih.gov/). Exposure time was constant between sections. The average intensity projection was analysed, taken from the central focal plane and 2 adjacent focal planes in either direction, with an interplane interval of 1 μm. The area of M1 ^61^ was measured using the freehand tool and cFos^+^ cells were manually counted. The average number of cFos^+^ cells/mm^2^ per mouse was calculated by dividing the mean number of counted cells by the area measured. Per animal 10-12 sections were counted.

### Statistical Analysis

Data was analysed using SPSS 23 (IBM Corporation, New York, USA), with an alpha level of p < 0.05 was regarded as significant throughout. All data was tested for normality using the Shapiro-Wilks test. Outliers were determined by both visual inspection of boxplots and/or standardised residuals of >±3 (https://statistics.laerd.com/). Unless otherwise stated analysis was conducted on untransformed data. Data was analysed by unpaired two-tailed *t-*test, Mann-Whitney U, one-way-ANOVA or mixed ANOVA. For *t-*tests that violate homogeneity of variances, Welch’s *t* test is used. For ANOVA where Mauchly’s assumption of sphericity was violated, the Greenhouse-Geisser correction was used. The assumptions of homogeneity of variances and covariances were assessed using Levene’s and Box’s M tests. Significant interactions were investigated with one-way ANOVAs.

## Results

### There was a significant reduction in *Dlg2* mRNA in the cortex of heterozygotic mice

There was no difference in the expression of *Dlg2* in the cerebellum (*t*_(9.301)_ = 0.547, *p* = 0.597, *t-*test) or hippocampus, (*t*_(21)_ = −0.238, *p* = 0.815, *t-*test) between WT and *Dlg2*^+/−^ mice (Figure 1a,b). However, there was a reduction in mRNA in the cortex in the *Dlg2*^+/−^ mice (*t*_(15)_ = −4.163, *p* = 0.001, *t-*test, Figure 1c). The expression of *Dlg1, Dlg3,* or *Dlg4* in the same brain regions was not altered in the *Dlg2*^+/−^ mice (*Supplementary Figure 1*).

**Figure 1.**
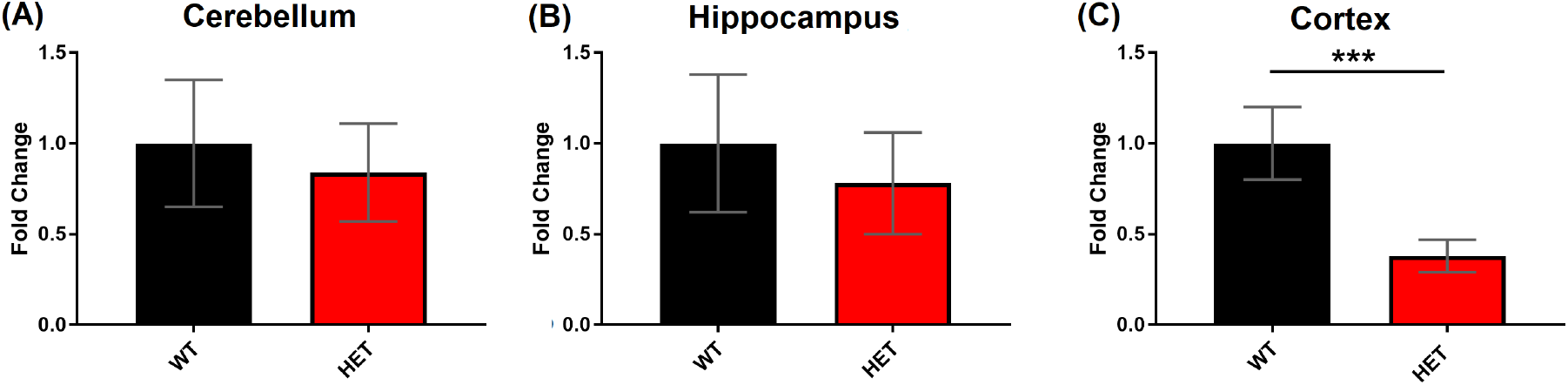
Region selective reduction in *Dlg2* expression in *Dlg2*^+/−^ mice. No changes in the expression of *Dlg2* mRNA in the cerebellum (a) or hippocampus (b) of *Dlg2*^+/−^ (HET) mice compared to WT were measured by RT-qPCR, however there was a reduction in the cortex (prefrontal cortex) (c). Error bars represent SEM. *n* = Cerebellum; 9 (WT), 8 (HET), hippocampus; 12 (WT), 11 (HET), and cortex: 8 (WT), 9 (HET). *** p < 0.001 (*t*-test).

### *Dlg2*^+/−^ mice exhibit impaired reactivity to an acoustic stimulus but intact PPI

The average of the first three acoustic stimulus pulses at 120 dB were analysed as an index of emotional reactivity. *Dlg2*^+/−^ mice exhibited a reduced reactivity to the 120 dB auditory stimulus (*t*_(40)_ = 3.303, *p* = 0.004, *t-*test) (Figure 2a). The reduced startle response in the *Dlg2*^+/−^ mice to the stimulus persisted across the first 6 pulse alone trials with no evidence of habituation (TRIAL: F _(3.825, 152,983)_ = 0.803, *p* = 0.520, GENOTYPE: F _(1, 40)_ = 7.441, *p* = 0.009, GENOTYPE × TRIAL: F _(3.825, 152.983)_ = 1.250, *p* = 0.293, mixed ANOVA) (Figure 2b).

**Figure 2.**
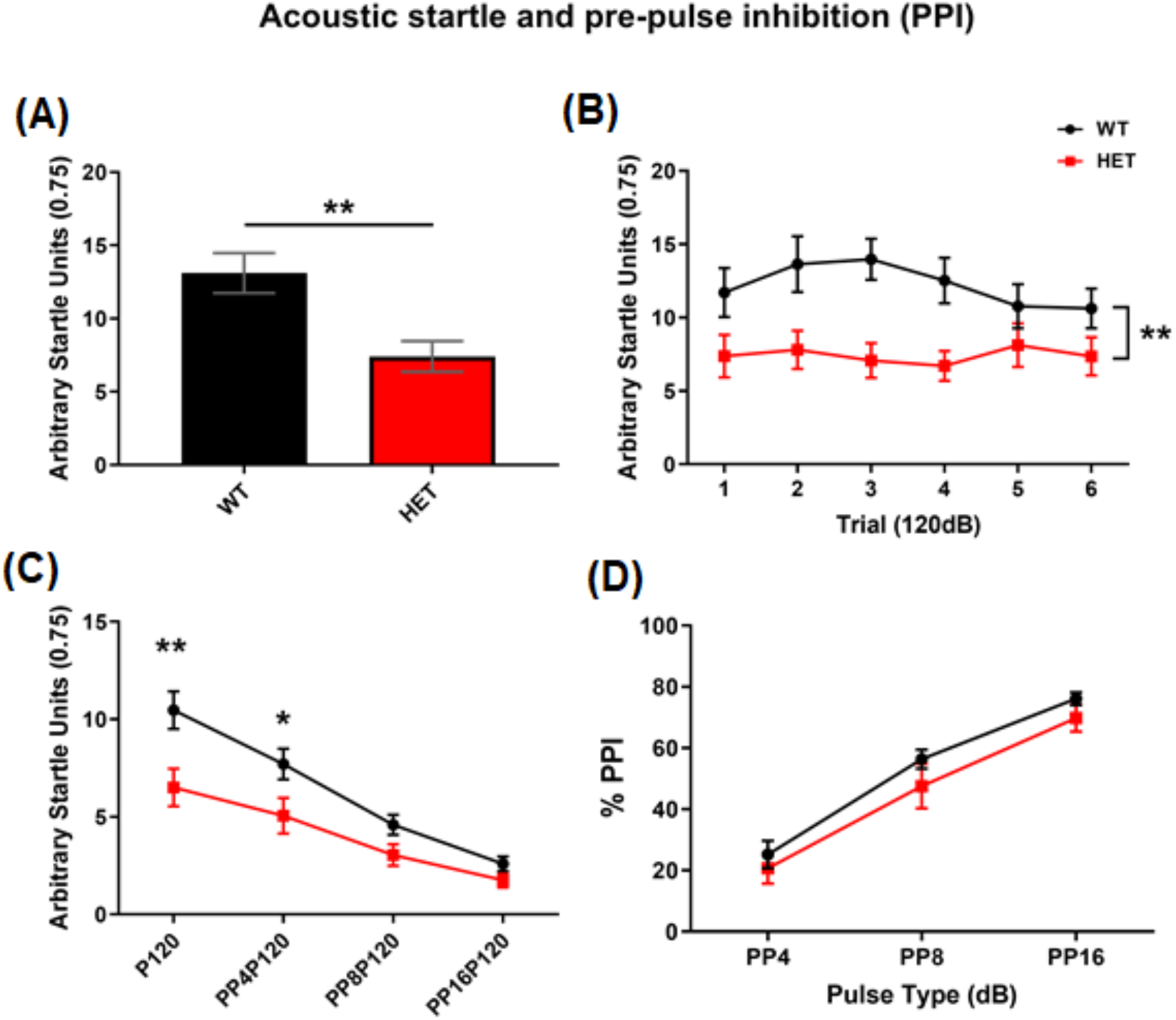
Reduced acoustic startle responses but intact PPI of acoustic startle in *Dlg2*^+/−^ mice. (a) *Dlg2*^+/−^ mice (HET) showed reduced emotional reactivity compared to WT as measured as the averaged startle response to the first three startle stimuli at 120 dB. (b) The difference in startle responses between the genotypes persisted across the first 6 pulses at 120 dB. (c) There was a monotonic reduction of the startle response by an increasing intensity prepulse stimulus 40 ms before the 120 dB startle stimulus in both WT and HET mice. This occurred despite a reduced startle response in the HETs without a pre-pulse (P120). (d) Both HET and WT mice exhibit a similar increase in startle response inhibition (% PPI) with pre-pulse intensity when expressed as a percentage of the 120 dB pulse-alone (P120). PP4P120, PP8P120, and PP16P120 = pre-pulse 4dB, 8dB and 16 dB above background noise (70dB), respectively. Error bars are SEM. n = 25 (WT) 17 (HET). HET compared to WT, * p < 0.05 (ANOVA), ** p < 0.01 (t-test (a), ANOVA (b) and (c)).

For PPI, as the intensity of the preceding pre-pulse increased the startle response decreased in both WT and *Dlg2*^+/−^ but with an apparent difference between the genotypes (PULSE: *F*_(1.742, 69.692)_ = 5.791, *p* = 0.007, GENOTYPE: *F*_(1, 40)_ = 5.916, *p* = 0.020, PULSE × GENOTYPE: *F*_(1.742, 69.692)_ = 5.791, *p* = 0.007, mixed ANOVA) (Figure 2c). The reduced responses in the *Dlg2*^+/−^ mice to the startle stimulus alone (P120) are likely to confound interpretation. When startle response was adjusted for the baseline response at 120 dB, there was no difference in the percentage PPI between the genotypes over the range of pre-pulse intensities tested (GENOTYPE: *F*_(1, 40)_ = 1.704, *p* = 0.199, PULSE: *F*_(1.730, 69.199)_ = 123.350, *p* = <0.01, GENOTYPE × PULSE: *F*_(1.730, 69.199)_ = 0.238, *p* = 0.757, mixed ANOVA) (Figure 2d). By using a lower intensity startle stimulus to ameliorate a potential confound of PPI measurement by the responses to the standard 120dB acoustic startle stimulus in the mutant animals, we showed that both genotypes showing similar emotional reactivity and PPI to a 105dB stimulus (*Supplementary Figure 2*). This suggests that failure to see an effect on absolute PPI in the *Dlg2*^+/−^ mice is not likely due to a ceiling or floor response effect masking modulation of startle responses. Thus, while the startle response to a loud acoustic stimuli in the *Dlg2*^+/−^ mice was reduced at least for a 120dB startle stimulus, sensorimotor gating was intact in *Dlg2*^+/−^ mice.

### *Dlg2*^+/−^ mice demonstrate impaired between-session habituation to a novel context

The daily total beam breaks for locomotor activity sessions to an initially novel open field (novel on Day 1) were compared between the genotypes using mixed ANOVA with the following factors: Genotype (WT and HET) and Day (1, 2, 3, 4, 5). Both genotypes exhibited reduced activity between sessions (DAY: *F*_(4, 132)_ = 31.229, *p* = < 0.001, GENOTYPE: *F*_(1, 33)_ = 0.229, *p* = 0.636, DAY × GENOTYPE: *F*_(4, 132)_ = 2.264, *p* = 0.066) (Figure 3a).

**Figure 3.**
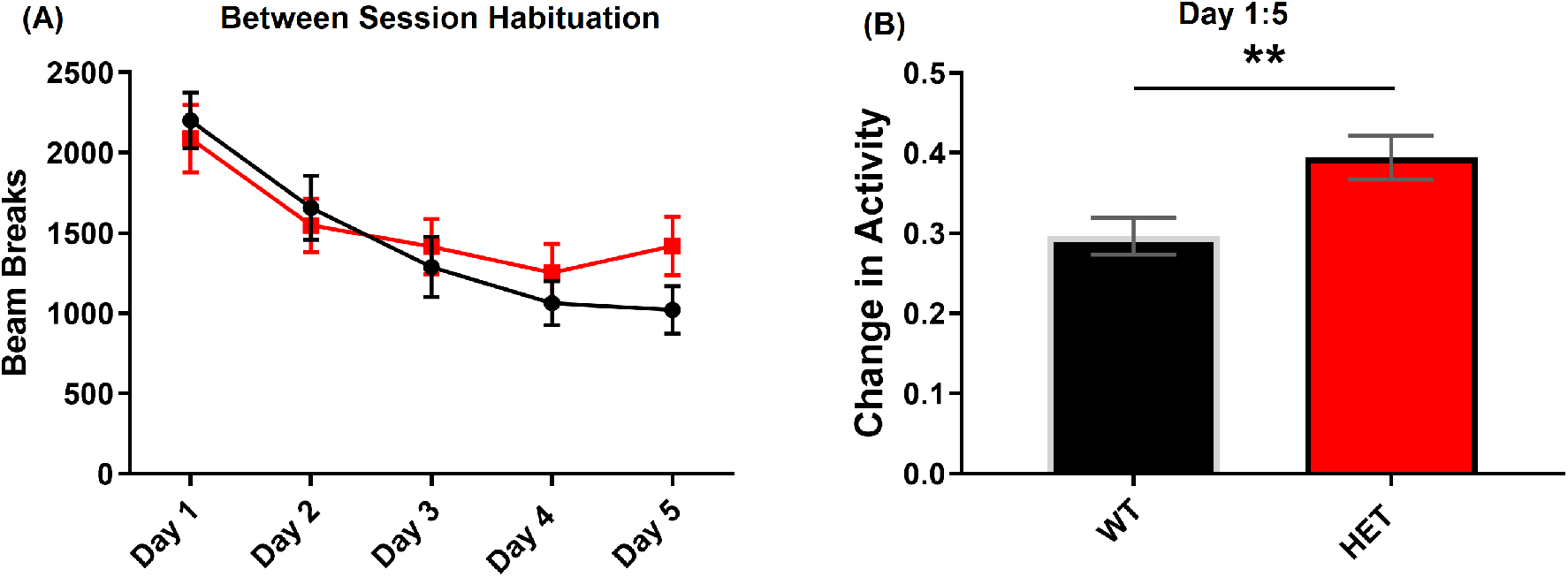
Impairment in the habituation to a novel context by *Dlg2*^+/−^ mice. (a) Both WT and *Dlg2*^+/−^ mice (HET) mice showed a similar decrease in total beam breaks for locomotor activity conducted in an activity box measured across 5 consecutive days. (b) When the change in activity between Day 1 and Day 5 was calculated (Day 5/(Day1 + Day 5)), *Dlg2*^+/−^ mice exhibited less change compared to WT mice, indicating weaker between-session habituation to the context after 5 days. Error bars are SEM. n = 18 (WT), 17 (HET). ** p = 0.01 (*t*-test).

The ratio of activity measured as between Day 1 to Day 5 was analysed as a measure of between-session habituation and compared between the genotypes using unpaired *t* test. *Dlg2*^+/−^ mice demonstrated less change in activity ratio at Day 5 and thus less habituation to the context than WT mice (*t*_(33)_ = −2.750, *p* = 0.010, *t*-test) (Figure 3b). No differences were found for within-session habituation on any day (*Supplementary Figure 3*). Therefore while the *Dlg2*^+/−^ mice appeared to show normal within-session habituation and memory of the context exposure, they showed weaker habituation after 5 days.

### *Dlg2*^+/−^ mice displayed impaired motor learning on two accelerating rotarod protocols

In an initial experiment (Cohort 1), motor learning and function (performance) were measured using an accelerating (Days 1 and 2) and fixed speed rotarod task (Days 3-8), respectively (Figure 4a). *Dlg2*^+/−^ mice exhibited an impairment on the accelerating rotarod, indicating deficient motor learning (TRIAL: *F*_(4, 168)_ = 35.318, *p* = <0.001, GENOTYPE: *F*_(1, 42)_ = 1.952, *p* = 0.170, TRIAL × GENOTYPE: F _(4, 168)_ = 3.598, *p* = 0.008, mixed ANOVA) (Figure 4b). However, motor function as assessed on a fixed speed rotarod, was intact in *Dlg2*^+/−^ mice (TRIAL: *F*_(5.696, 233.53)_ = 120.649, *p* = < 0.001, GENOTYPE: *F*_(1, 41)_ = 0.147, *p* = 0.704, TRIAL × GENOTYPE: *F*_(5.696, 233.53)_ = 0.631, *p* = 0.697, mixed ANOVA) (Figure 4c).

**Figure 4.**
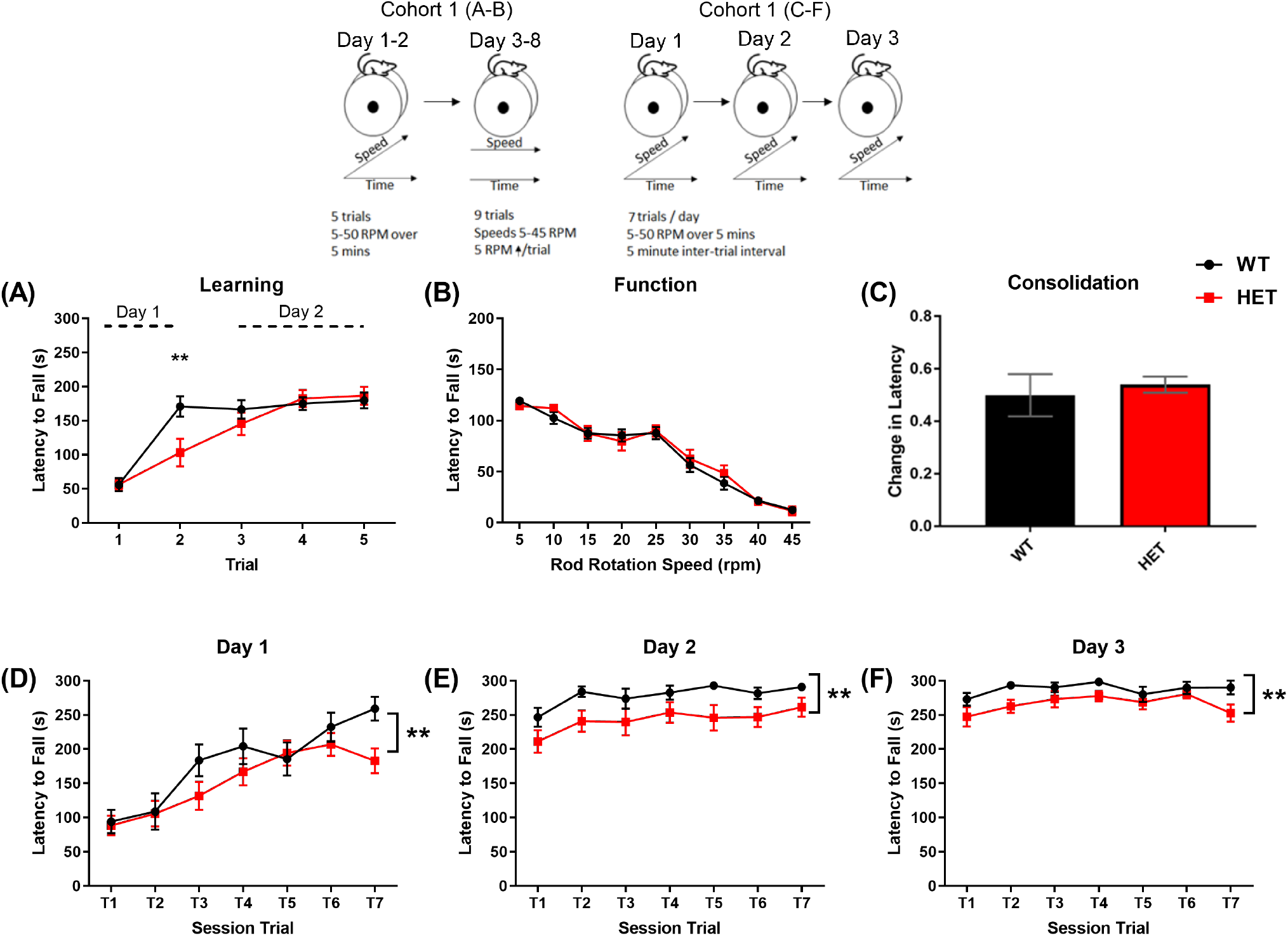
*Dlg2*^+/−^ mice exhibit slower motor learning but intact normal motor function and retention of motor memory across days. (a) Schematics of two independent rotarod experiments. Cohort 1 (*n* = 25 (WT), 18 (HET)) were first trained and tested using an accelerating rotarod protocol (Days 1 and 2) followed by a fixed speed protocol (Days 3-8) to assess motor learning and function, respectively. Motor learning in Cohort 2 was examined across three days using an accelerating rotarod protocol with more trials per day (7 trials) then the Cohort 1. (b) Cohort 1 *Dlg2*^+/−^ mice exhibited a shorter latency to fall than WT mice at Trial 2 only (*p* = 0.004) on the accelerating rotarod trials. The first 2 trials were conducted on Da remaining trials on Day 2 (c). In Cohort 1, the latency to fall for both genotypes decreased as the speed increased on the fixed rotarod task (Days 3-8). (d) In Cohort 2, both genotypes (*n* = 14 (WT), 16 (HET)) exhibit a similar decrease in latency to fall between the last trial of Day 1 and first trial of Day 2 (Day 2, Trial 1/(Day 1, Trial 7 + Day 2, Trial 1) indicating intact consolidation of the motor learning on Day 1. (e-g) Both genotypes in Cohort 2 demonstrated an increase in the latency to fall between trials on each of the three training days ((e) Day 1, *p* = < 0.001; (f) Day 2, *p* = 0.013; (g) Day 3, *p* = 0.030, AVOVA). However, *Dlg2*^+/−^ mice exhibited lower mean latencies to fall across trials than WT on all the three training days. Error bars represent SEM. ** *p* < 0.01 (ANOVA).

To further probe a motor learning deficit a more intensive accelerating rotarod protocol was employed, which allowed for investigation of fast and slow motor learning (Buitrago et al. 2004). An independent cohort of mice (Cohort 2) was trained using an accelerating rotarod task consisting of 7 trials/day for 3 consecutive days (Figure 4a). A TRIAL (*F*_(6, 168)_ = 16.713, *p* = <0.001, mixed ANOVA), DAY (*F*_(1.594, 44.630)_ = 147.891, *p* =<0.001, mixed ANOVA), DAY X TRIAL (*F* _(6.664, 186.591)_ = 7.255, *p* =<0.001, mixed ANOVA) interaction, but no DAY or TRIAL X GENOTYPE interactions (DAY X GENOTYPE: *F*_(1.594, 44.630)_ = 0.437, *p* = 0.604, TRIAL X GENOTYPE: *F*_(6, 168)_ = 0.711, *p* 0.626, *p* = <0.001, DAY X TRIAL X GENOTYPE: *F*_(6.664, 186.591)_ = 0.968, *p* = 0.454, mixed ANOVA) show that all mice show motor learning over the task (Figure 4e–g). Nevertheless, a GENOTYPE effect (*F*_(1, 28)_ = 12.059, *p* = 0.002, mixed ANOVA) and consistently lower latencies to fall measured in the *Dlg2*^+/−^ mice versus WT indicate weaker motor learning in heterozygous mice. In both genotypes motor performance on the last trial of Day 1 was retained on the first trial of Day 2, demonstrating that the consolidation of motor learning was unimpaired in *Dlg2*^+/−^ mice (*t*_(28)_ = −1.146, *p* = 0.261, *t*-test) (Figure 4d).

### *Dlg2*^+/−^ mice lack increased neuronal activity in M1 following motor learning

Using cFos expression as a proxy marker for neuronal activity ^56^, we investigated cFos levels using IHC in M1 of the motor cortex 90 minutes after completion of two trials of an accelerating rotarod task. The time point selected reflected the phase in training where we observed the largest differences in the latency to fall between the genotypes, and when peak activity-dependent cFos protein expression has been reported ^56,62^. M1 is a key cortical region associated with motor learning, in particular during the early phase of learning ^63^. Given the previously observed impairment in motor learning, we predicted that there would be less cFos expression as an index of reduced activity in M1 after training in the *Dlg2*^+/−^ mice.

During the motor skill learning task (Figure 5a), an increased latency to fall at Trial 2 compared to Trial 1 was observed (TRIAL: *F*_(1, 13)_ 6.602, *p* = 0.023, mixed ANOVA), but with no effect of GENOTYPE (F _(1, 13)_ = 0.215, *p* = 0.624, mixed ANOVA) or TRIAL X GENOTYPE interaction (*F*_(1, 13)_ = 1.602, *p* = 0.228, mixed ANOVA) apparent in this truncated learning protocol (Figure 5b).

**Figure 5.**
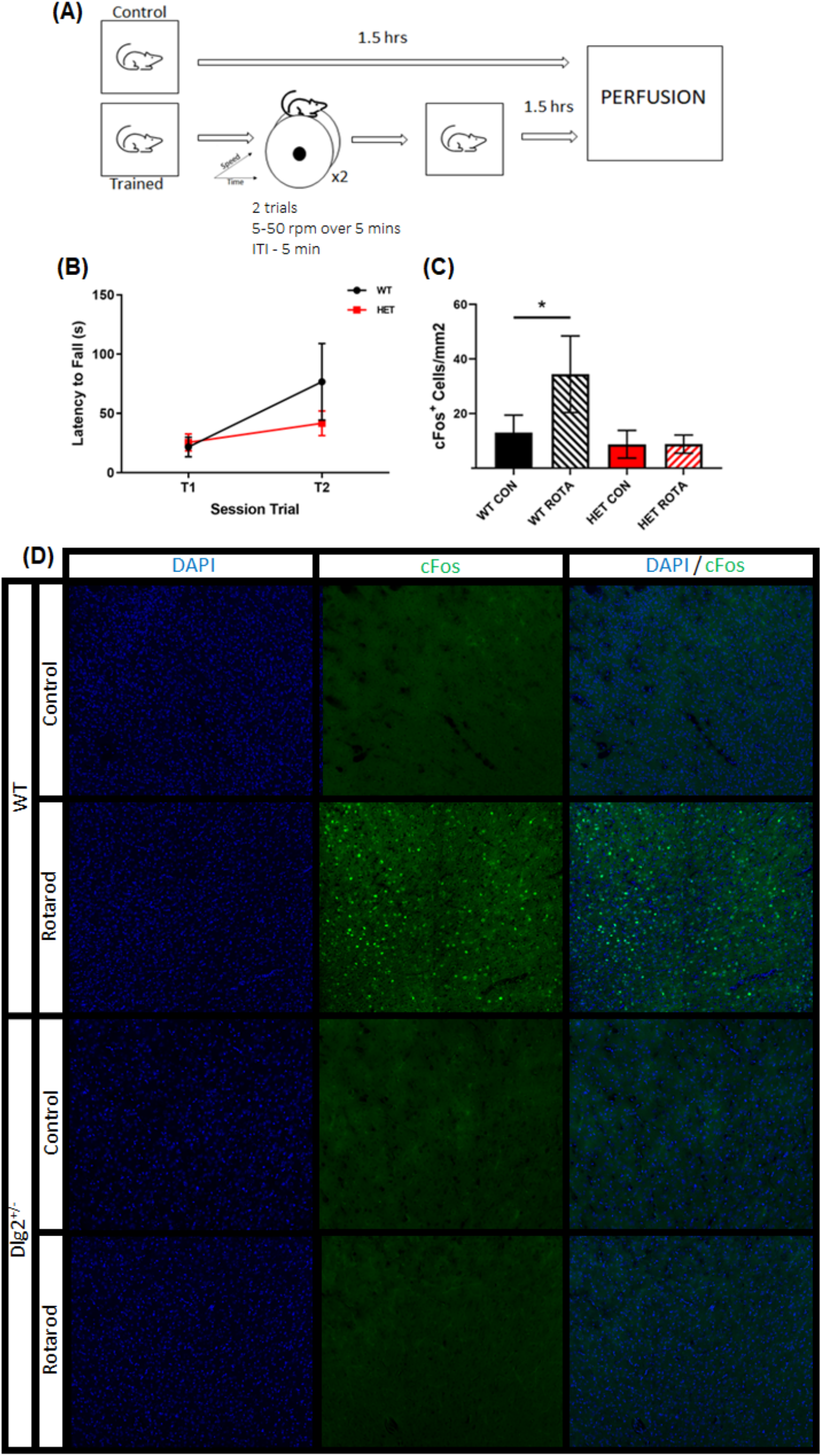
Reduced cFos expression after motor rod training in M1 of *Dlg2*^−/−^ mice. (a) Schematic of experimental design. Mice (ROTA) were single caged and underwent two trials on the accelerating rotarod protocol (0-50 rpm over 5 min, ITI = 5 min) then returned to cage for 1.5 hrs before perfusion. Control (CON) mice were single caged in the home cage for 1.5 hrs before perfusion. (b) Both WT and Dlg2^−/−^ (HET) mice show an increased latency to fall (s) between the first and second trial of the accelerating rotarod task (n = 7 (WT) 8 (HET)). (c) Only WT rotarod trained mice exhibited increased cFos expression in M1 90 min compared to single caged controls (n = 4 (WT CON), 6 (WT ROTA), 3 (HET CON), 7 (HET ROTA)). (d) Representative images of M1 stained for DAPI (blue) and cFos (green puncta). Error bars are SEM. * p < 0.01 (Mann Whitney-U).

The number of cFos positive neurons in M1 was increased in WT mice compared to home-cage naïve controls 90 minutes after Trial 2 (*U* = 12.000, *p* = 0.038, Mann Whitney-U test). No increase was observed in similarly trained mice *Dlg2*^+/−^ mice compared to home cage, *Dlg2*^+/−^ control mice (*U* = 38.00, *p* = 0.90, Mann Whitney-U test) (Figure 5c,d). Therefore while both genotypes show evidence of motor learning on the accelerating rotarod between Trials 1 and 2, an increase in cFos expression in M1 motor cortex was seen in the trained WT but not *Dlg2*^+/−^ mice as predicted.

## Discussion

We report that in a mouse model of Dlg2 heterozygosity, of high clinical relevance to psychiatric and developmental disorders ^32,64^, reduced expression of *Dlg2,* but not other Dlg family members, was observed in the cortex but not hippocampus or cerebellum. The model also exhibited impairments in motor learning and long-term habituation to a novel context, but not PPI of acoustic startle. The behavioural impairments were unconfounded by changes in anxiety as measured on the elevated plus maze or open field (*Supplementary Figure 4*). The impact of reduced genetic dosage on specific behaviours may reflect regional haploinsufficiency and thus localised reductions in function as evidenced by the reduced activity-dependent cFos expression in M1 motor cortex during motor learning. These impairments implicate Dlg2 in phenotypes associated with psychiatric and developmental disorders ^47–51,53–55,65,66^.

An impairment in motor skill learning but not motor co-ordination in *Dlg2*^+/−^ mice was shown in independent cohorts using two different rotarod training protocols. Using a sparse training protocol consisting of short training trials on an accelerating rotarod spread over 2 days, the *Dlg2*^+/−^ mice showed a markedly lower improvement in performance between the first two trials on Day 1 as shown by a smaller increase in the latency to fall compared to WT. In WT, the greatest improvement in performance is seen between trials 1 and 2 on Day 1 and is consistent with the typical pattern seen with learning ^67^. Nevertheless the performance of the heterozygous mutants matched those in WT during subsequent trials on Day 2. The similarity in performance by the end of training and a formal assessment of motor function on the fixed speed rotarod after motor skill acquisition show that motor function in the *Dlg2*^+/−^ mice was unaffected *per se,* and that the motor learning deficit observed was not confounded by basic deficits in motor function. The pattern of deficit in the period of rapid learning observed on the first day, equating with within-session training, reflects an impact of Dlg2 heterozygosity on the “fast” phase of motor skill learning ^63,67,68^. Smaller performance improvements seen in the *Dlg2*^+/−^ mice in early trials might also be explained by changes including altered attention, motivation or physiological “warm up” effects that are more specifically associated with within-session ^69–72^. However, using a more prolonged motor learning protocol (7 trials/d for 3 d) to better observe learning *and* memory function, *Dlg2*^+/−^ mice similarly showed deficits in early within-session learning. But they also showed a performance deficit that persisted across at least three days of training. The mutants showed retention of performance gains between the last trial on Day 1 and the first on Day 2 indicating that between-session consolidation of motor skill memory was intact in the *Dlg2*^+/−^ mice. Thus overall, *Dlg2*^+/−^ mice show deficits in both “fast” within-session and “slow” between– session motor skill learning, but intact motor function and memory consolidation.

The within-session motor learning was associated with an increase in cFos expression in M1 motor cortex in WT mice. The expression of cFos is commonly used as molecular marker of synaptic plasticity and learning ^73–76^. The observed increase in expression demonstrates the activation M1 by the rotarod protocols we employed, which is consistent with the critical contribution of M1 to fast motor skill learning in rodents ^63^ and to fast and slow motor skill learning in both rodents and humans ^77^. The failure to see a similar increase in cFos expression in the *Dlg2*^+/−^ mice correlates with their impairment in motor learning and implicate DLG2-associated plasticity mechanisms, particularly long-term potentiation (LTP) ^41^ to support motor learning.

While the plasticity processes in motor cortex are associated with fast and slow learning, the loci and the contributing circuits, including subcortical structures, are distinct ^63,77,78^ and likely involve the selective engagement of neuronal subtypes or clusters ^79^. Furthermore, the activated networks that represent the motor memory “engram” continue to be refined through both within- and between-session motor skill learning. The deficits in both fast and slow motor skill learning in the *Dlg2*^+/−^ mice are consistent with a broad role for DLG2 in motor leaning-associated plasticity processes in M1 (this study) and associated circuits more widely ^40^. While the discrepancy we see in our two learning experiments regarding between-session learning, with *Dlg2*^+/−^ mice showing persistently lower performance between sessions only when trained using longer sessions over several days, cannot be explained by a difference in the task used (they are the same), it is possible that prolonged training engages different circuits such as cortico-striatal pathways involved in developing habitual responses ^78^. We speculate that these circuits may also be more sensitive to the reduced DLG2 dosage in this heterozygous mouse model.

Deficits in between-session habituation to a novel context in the *Dlg2*^+/−^ mice manifested as a smaller decrease in locomotor activity after five consecutive days than WT. Repeated exposure to an unfamiliar stimulus, such as a context, results in reduced responses, including locomotor activity, over time as the stimulus becomes less novel ^80,81^. Nevertheless, within-session habituation was similar in both genotypes, perhaps reflecting a greater impact of DLG2 heterozygosity on long-term than short-term habituation and the different molecular and plasticity mechanisms supporting them ^82^. Notably long-term habituation requires protein synthesis for its production and maintenance ^80,83^ similar to NMDA receptor-mediated plasticity mechanisms regulating long-term associative memory ^84^. Interestingly, habituation impairments have been reported in *Gria1* knockout mice ^85^. The authors proposed that deficits in habituation are central to psychological processes such as sensitization and aberrant salience that give rise to psychosis associated with schizophrenia and other psychiatric conditions. Together with our data, this suggests disruptions that AMPAR-mediated signalling involving DLG2 at glutamatergic synapses is a mechanism underlying behavioural habituation deficits that are core for psychosis.

To our knowledge this was the first study investigating acoustic startle and PPI in *Dlg2* mutants. PPI deficits are a common feature of a range of neuropsychiatric disorders serving as an endophenotype to identify risk genes in humans and to illuminate neural processes and circuits affected in genetic models in animals ^86^. The *Dlg2*^+/−^ mice show no impairment in PPI. A decrease in reactivity to the standard 120 dB startle stimulus in the mutants is unlikely to account for the inability to detect a change in PPI because PPI was similarly unaffected using a 105dB intensity startle stimulus where reactivity in the two genotypes was matched. The decreased startle reactivity is unlikely to be due to a gross motor deficit as motor co-ordination and initial locomotor activity an open field were unaffected in the *Dlg2*^+/−^ mice. Thus, whilst an explanation regarding the impaired acoustic startle responses needs to be elucidated, reduced dosage of Dlg2 does not appear to affect sensorimotor gating in this mouse model.

Our observations of unaffected motor performance and locomotor activity in the *Dlg2*^+/−^ mice can be viewed as broadly concordant with previous investigations using *DLG2* models when considering gene dosage. Motor coordination and performance in the *Dlg2*^+/−^ deletion model has been reported to be either unaffected ^87^ or mildly impaired, but with the impairment absent was in the heterozygous mice 39. Both *Dlg2*^+/− 42^ and *Dlg2* ^⍰E14/⍰⍰E14^ (the same exon targeted as in the mice used in our study) deletion models show mild hypoactivity activity in novel open fields but not in familiar environments ^40^. Together the data suggest that while subtle effects can be seen on motor function-related behaviours in deletion models, these deficits are absent when one copy of *DLG2* is present. The dosage effect may highlight a role for DLG2 in locomotor-related behaviour, including locomotor behaviours associated with novelty detection, but the role is redundant because DLG paralogs are able to compensate for a reduction of DLG2 function. This hypothesis may equally apply to another behavioural phenotype observed in genetic *DLG2* models, that of altered social behaviours which is seen in DLG homozygotes ^39,40^ but not heterozygotes ^39^. Thus, the utility of heterozygous models is not only in interrogating more accurate genetic risk models for human disease but also for identifying the essential, or selective functions, associated with the targeted gene. Additionally, it is important to note that we measured reduced expression of DLG2 in some brain regions (cortical) but not others (hippocampus and cerebellum). A differential impact on regional expression levels if recapitulated more globally may be another source of specificity and variability in observed functional deficits within and between genetic models.

The selectively of the impairments we see in Dlg2^+/−^ mice may reflect the impact of DLG haploinsufficiency on long-term synaptic plasticity mechanisms within functional circuitries. Deficits in long-term motor learning and habitation may be indicative of impairments in synaptic plasticity mechanisms in the cortico-striato-thalamo-cortical and cortico-cerebello-thalamo-cortico pathways underlying motor learning (as discussed earlier), and the medial temporal-midbrain-striatal pathways that mediate the novelty-to-familiarly transition of stimuli that manifest as habituation ^88^. DLG2 is critical for long-term potentiation (LTP), which underlies long-term protein-synthesis dependent learning, but not AMPA and NMDA receptor–mediated synaptic transmission ^41^ that are extant for information processing and behavioural responses not dependent on LTP such as motor co-ordination, PPI, and anxiety. In comparison to DLG2 mutants, DLG4 (PSD-95) models show alterations in both glutamatergic long-term plasticity and basal transmission ^41^ and also more severe and wider ranging behavioural affects ^42^. These observations lend further support for a role for DLG2 in functions specifically dependent on long-term plasticity.

## Conclusions

In this paper, we present data indicating heterozygosity of *Dlg2,* which has been previously associated with a number of psychiatric diseases, results in selective behavioural deficits similarly associated with disease in *Dlg2*^+/−^ mice. The deficits in motor learning and long-term habituation of locomotor responses to a novel context that we observed in the mutant mice can broadly be considered as failures to adapt to the environment. These are also functions dependent on synaptic plasticity mechanisms, which DLG2 mediates. The reduced cortical regulation of the plasticity-associated gene cFos during motor learning may indeed be a molecular signature for impairments in synaptic plasticity in DLG2 mutant models. Other functions including sensorimotor gating of an acoustic stimulus measured by PPI, motor co-ordination and anxiety were unaffected in this *Dlg2*^+/−^ mouse model. This suggests functional behavioural and cognitive domains associated with long-term adaptation to the environment that also depend on specific long-term plasticity mechanisms are particularly sensitive to haploinsufficiency of DLG2 across psychiatric disorders.

## Supporting information

051021 Pass Supplemental Information

## Acknowledgements

This work was also supported by a Wellcome Trust Strategic Award ‘DEFINE’ grant no. 100202/Z/12/Z and core support from the Neuroscience and Mental Health Research Institute, Cardiff University, UK.

## References

1. Rapoport J, Giedd J, Gogtay N. Neurodevelopmental model of schizophrenia: update 2012. Mol Psychiatry. 2012;17(12):1228. doi:10.1038/MP.2012.23

2. Arlington V. Diagnostic and Statistical Manual of Mental Disorders. 5th ed. American Psychiatric Association; 2013.

3. Reiss AL. Childhood developmental disorders: an academic and clinical convergence point for psychiatry, neurology, psychology and pediatrics. J Child Psychol Psychiatry. 2009;50(1–2):87–98. doi:10.1111/J.1469-7610.2008.02046.X

4. Moreno-De-Luca D, Sanders SJ, Willsey AJ, et al. Using large clinical data sets to infer pathogenicity for rare copy number variants in autism cohorts. Mol Psychiatry. 2013;18(10):1090–1095. doi:10.1038/mp.2012.138

5. Capute, Accardo P. The Spectrum of Neurodevelopmental Disabilities, Capute & Accardo’s Neurodevelopmental Disabilities in Infancy and Childhood: Neurodevelopmental Diagnosis and Treatment. 3rd Vol 2. Brookes Publishing; 2008.

6. Kirov G, Pocklington AJ, Holmans P, et al. De novo CNV analysis implicates specific abnormalities of postsynaptic signalling complexes in the pathogenesis of schizophrenia. Mol Psychiatry. 2012;17(2):142–153. doi:10.1038/mp.2011.154

7. Anttila V, B B-S, HK F, et al. Analysis of shared heritability in common disorders of the brain. Science. 2018;360(6395). doi:10.1126/SCIENCE.AAP8757

8. Gandal MJ, Haney JR, Parikshak NN, et al. Shared molecular neuropathology across major psychiatric disorders parallels polygenic overlap. Science (80-). 2018;359(6376):693–697. doi:10.1126/SCIENCE.AAD6469

9. Singh T, Walters JTR, Johnstone M, et al. The contribution of rare variants to risk of schizophrenia in individuals with and without intellectual disability. Nat Genet. 2017;49(8):1167–1173. doi:10.1038/ng.3903

10. Grove J, Ripke S, Als TD, et al. Identification of common genetic risk variants for autism spectrum disorder. Nat Genet. 2019;51(3):431–444. doi:10.1038/s41588-019-0344-8

11. Doherty JL, Owen MJ. Genomic insights into the overlap between psychiatric disorders: implications for research and clinical practice. Genome Med. 2014;6(4):29. doi:10.1186/gm546

12. Pardiñas AF, Holmans P, Pocklington AJ, et al. Common schizophrenia alleles are enriched in mutation-intolerant genes and in regions under strong background selection. Nat Genet. 2018;50(3):381–389. doi:10.1038/s41588-018-0059-2

13. Pocklington AJ, O’Donovan M, Owen MJ. The synapse in schizophrenia. Eur J Neurosci. 2014;39(7):1059–1067. doi:10.1111/ejn.12489

14. Ripke S, Neale BM, Corvin A, et al. Biological insights from 108 schizophrenia-associated genetic loci. Nature. 2014;511:421–427. doi:10.1038/nature13595

15. Fromer M, Pocklington AJ, Kavanagh DH, et al. De novo mutations in schizophrenia implicate synaptic networks. Nature. 2014;506(7487):179–184. doi:10.1038/nature12929

16. Glessner JT, Reilly MP, Kim CE, et al. Strong synaptic transmission impact by copy number variations in schizophrenia. Proc Natl Acad Sci. 2010;107(23):10584–10589. doi:10.1073/pnas.1000274107

17. Network and Pathway Analysis Subgroup of Psychiatric Genomics Consortium TN and PAS of the PG. Psychiatric genome-wide association study analyses implicate neuronal, immune and histone pathways. Nat Neurosci. 2015;18(2):199–209. doi:10.1038/nn.3922

18. van de Lagemaat LN, Grant SGN. Genome Variation and Complexity in the Autism Spectrum. Neuron. 2010;67(1):8–10. doi:10.1016/J.NEURON.2010.06.026

19. Chung BH-Y, Tao VQ, Tso WW-Y. Copy number variation and autism: New insights and clinical implications. J Formos Med Assoc. 2014;113(7):400–408. doi:10.1016/J.JFMA.2013.01.005

20. Leblond CS, Nava C, Polge A, et al. Meta-analysis of SHANK Mutations in Autism Spectrum Disorders: A Gradient of Severity in Cognitive Impairments. Barsh GS, ed. PLoS Genet. 2014;10(9):e1004580. doi:10.1371/journal.pgen.1004580

21. Voineagu I, Wang X, Johnston P, et al. Transcriptomic analysis of autistic brain reveals convergent molecular pathology. Nat 2011 4747351. 2011;474(7351):380–384. doi:10.1038/nature10110

22. Guang S, Pang N, Deng X, et al. Synaptopathology Involved in Autism Spectrum Disorder. Front Cell Neurosci. 2018;12:470. doi:10.3389/FNCEL.2018.00470

23. Gilman SR, Iossifov I, Levy D, Ronemus M, Wigler M, Vitkup D. Rare de novo variants associated with autism implicate a large functional network of genes involved in formation and function of synapses. Neuron. 2011;70(5):898–907. doi:10.1016/j.neuron.2011.05.021

24. Krumm N, O’Roak BJ, Shendure J, Eichler EE. A de novo convergence of autism genetics and molecular neuroscience. Trends Neurosci. 2014;37(2):95–105. doi:10.1016/j.tins.2013.11.005

25. Zhu J, Shang Y, Zhang M. Mechanistic basis of MAGUK-organized complexes in synaptic development and signalling. Nat Rev Neurosci. 2016;17(4):209–223. doi:10.1038/nrn.2016.18

26. Won S, Levy JM, Nicoll RA, Roche KW. MAGUKs: multifaceted synaptic organizers. Curr Opin Neurobiol. 2017;43:94–101. doi:10.1016/j.conb.2017.01.006

27. Walsh T, McClellan JM, McCarthy SE, et al. Rare Structural Variants Disrupt Multiple Genes in Neurodevelopmental Pathways in Schizophrenia. Science (80-). 2008;320(5875). Accessed April 18, 2017. http://science.sciencemag.org/content/320/5875/539.full

28. Xu B, Roos JL, Levy S, van Rensburg EJ, Gogos JA, Karayiorgou M. Strong association of de novo copy number mutations with sporadic schizophrenia. Nat Genet. 2008;40(7):880–885. doi:10.1038/ng.162

29. Purcell SM, Moran JL, Fromer M, et al. A polygenic burden of rare disruptive mutations in schizophrenia. Nature. 2014;506(7487):185–190. doi:10.1038/nature12975

30. Egger G, Roetzer KM, Noor A, et al. Identification of risk genes for autism spectrum disorder through copy number variation analysis in Austrian families. Neurogenetics. 2014;15(2):117–127. doi:10.1007/s10048-014-0394-0

31. Xing J, Kimura H, Wang C, et al. Resequencing and Association Analysis of Six PSD-95-Related Genes as Possible Susceptibility Genes for Schizophrenia and Autism Spectrum Disorders. Sci Rep. 2016;6:27491. doi:10.1038/srep27491

32. Ruzzo EK, Pérez-Cano L, Jung JY, et al. Inherited and De Novo Genetic Risk for Autism Impacts Shared Networks. Cell. 2019;178(4):850–866.e26. doi:10.1016/j.cell.2019.07.015

33. Noor A, Lionel AC, Cohen-Woods S, et al. Copy number variant study of bipolar disorder in Canadian and UK populations implicates synaptic genes. Am J Med Genet Part B Neuropsychiatr Genet. 2014;165(4):303–313. doi:10.1002/ajmg.b.32232

34. Harrison PJ, Weinberger DR. Schizophrenia genes, gene expression and neuropathology: on the matter of their convergence. Mol Psychiatry. 2005;10(1):40–68. doi:10.1038/sj.mp.4001558

35. Frantseva M V., Fitzgerald PB, Chen R, Moller B, Daigle M, Daskalakis ZJ. Evidence for Impaired Long-Term Potentiation in Schizophrenia and Its Relationship to Motor Skill Leaning. Cereb Cortex. 2008;18(5):990–996. doi:10.1093/cercor/bhm151

36. Chen CH, Huang CC, Cheng MC, et al. Genetic analysis of GABRB3 as a candidate gene of autism spectrum disorders. Mol Autism. 2014;5(1). doi:10.1186/2040-2392-5-36

37. Bourgeron T. From the genetic architecture to synaptic plasticity in autism spectrum disorder. Nat Rev Neurosci. 2015;16(9):551–563. doi:10.1038/nrn3992

38. Nithianantharajah J, Komiyama NH, McKechanie A, et al. Synaptic scaffold evolution generated components of vertebrate cognitive complexity. Nat Neurosci. 2013;16(1):16–24. doi:10.1038/nn.3276

39. Winkler D, Daher F, Wüstefeld L, et al. Hypersocial behavior and biological redundancy in mice with reduced expression of PSD95 or PSD93. Behav Brain Res. 2018;352:35–45. doi:10.1016/j.bbr.2017.02.011

40. Yoo T, Kim SG, Yang SH, Kim H, Kim E, Kim SY. A DLG2 deficiency in mice leads to reduced sociability and increased repetitive behavior accompanied by aberrant synaptic transmission in the dorsal striatum. Mol Autism. 2020;11(1):19. doi:10.1186/s13229-020-00324-7

41. Carlisle HJ, Fink AE, Grant SGN, O’Dell TJ. Opposing effects of PSD-93 and PSD-95 on long-term potentiation and spike timing-dependent plasticity. J Physiol. 2008;586(Pt 24):5885–5900. doi:10.1113/jphysiol.2008.163469

42. Nithianantharajah J, Komiyama NH, McKechanie A, et al. Synaptic scaffold evolution generated components of vertebrate cognitive complexity. Nat Neurosci. 2012;16(1):16–24. doi:10.1038/nn.3276

43. Favaro PD, Huang X, Hosang L, et al. An opposing function of paralogs in balancing developmental synapse maturation. PLoS Biol. 2018;16(12):e2006838. doi:10.1371/journal.pbio.2006838

44. Kirov G, Pocklington AJ, Holmans P, et al. De novo CNV analysis implicates specific abnormalities of postsynaptic signalling complexes in the pathogenesis of schizophrenia. Mol Psychiatry. 2012;17(2):142–153. doi:10.1038/mp.2011.154

45. Reggiani C, Coppens S, Sekhara T, et al. Novel promoters and coding first exons in DLG2 linked to developmental disorders and intellectual disability. Genome Med. 2017;9(1):67. doi:10.1186/s13073-017-0452-y

46. Braff DL, Grillon C, Geyer MA. Gating and habituation of the startle reflex in schizophrenic patients. Arch Gen Psychiatry. 1992;49(3):206–215. Accessed March 3, 2019. http://www.ncbi.nlm.nih.gov/pubmed/1567275

47. Gottesman II, Gould TD. The Endophenotype Concept in Psychiatry: Etymology and Strategic Intentions. Am J Psychiatry. 2003;160(4):636–645. doi:10.1176/appi.ajp.160.4.636

48. Walters JTR, Owen MJ. Endophenotypes in psychiatric genetics. Mol Psychiatry. 2007;12(10):886–890. doi:10.1038/sj.mp.4002068

49. Mena A, Ruiz-Salas JC, Puentes A, Dorado I, Ruiz-Veguilla M, De la Casa LG. Reduced Prepulse Inhibition as a Biomarker of Schizophrenia. Front Behav Neurosci. 2016;10:202. doi:10.3389/fnbeh.2016.00202

50. Kohl S, Wolters C, Gruendler TOJ, Vogeley K, Klosterkötter J, Kuhn J. Prepulse inhibition of the acoustic startle reflex in high functioning autism. PLoS One. 2014;9(3):e92372. doi:10.1371/journal.pone.0092372

51. Perry W, Minassian A, Lopez B, Maron L, Lincoln A. Sensorimotor Gating Deficits in Adults with Autism. Biol Psychiatry. 2007;61(4):482–486. doi:10.1016/j.biopsych.2005.09.025

52. Swerdlow NR, Braff DL, Geyer MA. Sensorimotor gating of the startle reflex: what we said 25 years ago, what has happened since then, and what comes next. J Psychopharmacol. 2016;30(11):1072. doi:10.1177/0269881116661075

53. Matsuo J, Ota M, Hidese S, et al. Sensorimotor Gating in Depressed and Euthymic Patients with Bipolar Disorder: Analysis on Prepulse Inhibition of Acoustic Startle Response Stratified by Gender and State. Front psychiatry. 2018;9:123. doi:10.3389/fpsyt.2018.00123

54. McDiarmid TA, Bernardos AC, Rankin CH. Habituation is altered in neuropsychiatric disorders—A comprehensive review with recommendations for experimental design and analysis. Neurosci Biobehav Rev. 2017;80:286–305. doi:10.1016/j.neubiorev.2017.05.028

55. MJ C, E G de J, MS C, et al. Motor abnormalities in first-episode psychosis patients and long-term psychosocial functioning. Schizophr Res. 2018;200:97–103. doi:10.1016/J.SCHRES.2017.08.050

56. Kovács KJ. Measurement of Immediate-Early Gene Activation-*c-fos* and Beyond. J Neuroendocrinol. 2008;20(6):665–672. doi:10.1111/j.1365-2826.2008.01734.x

57. Kleiber M. Body size and metabolism. Hilgardia. 1932;6:315–351.

58. Geyer MA, Dulawa SC. Assessment of Murine Startle Reactivity, Prepulse Inhibition, and Habituation. Curr Protoc Neurosci. Published online 2003:1–15. doi:10.1002/0471142301.ns0817s24

59. Elias GM, Funke L, Stein V, Grant SG, Bredt DS, Nicoll RA. Synapse-Specific and Developmentally Regulated Targeting of AMPA Receptors by a Family of MAGUK Scaffolding Proteins. Neuron. 2006;52:307–320. doi:10.1016/j.neuron.2006.09.012

60. Schmittgen TD, Livak KJ. Analyzing real-time PCR data by the comparative CT method. Nat Protoc. 2008;3(6):1101–1108. doi:10.1038/nprot.2008.73

61. Franklin KBJ, Paxinos G. Paxinos and Franklin’s The Mouse Brain in Stereotaxic Coordinates.

62. W T, R G. Activation of immediate early genes and memory formation. Cell Mol Life Sci. 1999;55(4):564–574. doi:10.1007/S000180050315

63. Costa RM, Cohen D, Nicolelis MAL. Differential corticostriatal plasticity during fast and slow motor skill learning in mice. Curr Biol. 2004;14(13):1124–1134. doi:10.1016/j.cub.2004.06.053

64. Reggiani C, Coppens S, Sekhara T, et al. Novel promoters and coding first exons in DLG2 linked to developmental disorders and intellectual disability. Genome Med. 2017;9(1):67. doi:10.1186/s13073-017-0452-y

65. Kemner C, Oranje B, Verbaten MN, van Engeland H. Normal P50 gating in children with autism. J Clin Psychiatry. 2002;63(3):214–217. Accessed February 11, 2019. http://www.ncbi.nlm.nih.gov/pubmed/11926720

66. Perry W, Minassian A, Feifel D, Braff DL. Sensorimotor gating deficits in bipolar disorder patients with acute psychotic mania. Biol Psychiatry. 2001;50(6):418–424. Accessed February 11, 2019. http://www.ncbi.nlm.nih.gov/pubmed/11566158

67. Karni A, Meyer G, Rey-Hipolito C, et al. The acquisition of skilled motor performance: fast and slow experience-driven changes in primary motor cortex. Proc Natl Acad Sci U S A. 1998;95(3):861–868. doi:10.1073/PNAS.95.3.861

68. Luft AR, Buitrago MM. Stages of Motor Skill Learning. Mol Neurobiol. 2005;32(3):205–216. doi:10.1385/MN:32:3:205

69. Mosberger AC, Clauser L de, Kasper H, Schwab ME. Motivational state, reward value, and Pavlovian cues differentially affect skilled forelimb grasping in rats. Learn Mem. 2016;23(6):289–302. doi:10.1101/LM.039537.115

70. Wulf G. Attention and Motor Skill Learning. Human Kinetics; 2007. Accessed October 5, 2021. https://psycnet.apa.org/record/2007-04641-000

71. Hoffman J. Warm-up and flexibility. In: Physiological Aspects of Sport Training and Performance. Human Kinetics; 2002:155–168.

72. Haibach PR, Reid G CD. Motor Learning and Development. In: Human Kinetics., ed.; 2011.

73. Curran T, Morgan JI. Memories of fos. BioEssays. 1987;7(6):255–258. doi:10.1002/BIES.950070606

74. Goelet P, Castellucci VF, Schacher S, Kandel ER. The long and the short of long–term memory—a molecular framework. Nat 1986 3226078. 1986;322(6078):419–422. doi:10.1038/322419a0

75. Grimm R, Schicknick H, Riede I, et al. Suppression of c-fos induction in rat brain impairs retention of a brightness discrimination reaction. Learn Mem. 1997;3(5):402–413. doi:10.1101/LM.3.5.402

76. de Hoz L, Gierej D, Lioudyno V, et al. Blocking c-Fos Expression Reveals the Role of Auditory Cortex Plasticity in Sound Frequency Discrimination Learning. Cereb Cortex. 2018;28(5):1645–1655. doi:10.1093/CERCOR/BHX060

77. Dayan E, Cohen LG. Neuroplasticity Subserving Motor Skill Learning. Neuron. 2011;72(3):443–454. doi:10.1016/J.NEURON.2011.10.008

78. Yin HH, Mulcare SP, Hilário MRF, et al. Dynamic reorganization of striatal circuits during the acquisition and consolidation of a skill. Nat Neurosci. 2009;12(3):333–341. doi:10.1038/nn.2261

79. Papale AE, Hooks BM. Circuit changes in motor cortex during motor skill learning. Neuroscience. 2018;368:283–297. doi:10.1016/j.neuroscience.2017.09.010

80. Rankin CH, Abrams T, Barry RJ, et al. Habituation revisited: an updated and revised description of the behavioral characteristics of habituation. Neurobiol Learn Mem. 2009;92(2):135–138. doi:10.1016/j.nlm.2008.09.012

81. Leussis MP, Bolivar VJ. Habituation in rodents: A review of behavior, neurobiology, and genetics. Neurosci Biobehav Rev. 2006;30(7):1045–1064. doi:10.1016/J.NEUBIOREV.2006.03.006

82. McDiarmid TA, Belmadani M, Liang J, et al. Systematic phenomics analysis of autism-associated genes reveals parallel networks underlying reversible impairments in habituation. Proc Natl Acad Sci. 2020;117(1):656–667. doi:10.1073/PNAS.1912049116

83. Ramaswami M. Network Plasticity in Adaptive Filtering and Behavioral Habituation. Neuron. 2014;82(6):1216–1229. doi:10.1016/J.NEURON.2014.04.035

84. Sweatt JD. Neural plasticity and behavior – sixty years of conceptual advances. J Neurochem. 2016;139:179–199. doi:10.1111/JNC.13580

85. Barkus C, Sanderson DJ, Rawlins JNP, Walton ME, Harrison PJ, Bannerman DM. What causes aberrant salience in schizophrenia? A role for impaired short-term habituation and the GRIA1 (GluA1) AMPA receptor subunit. Mol Psychiatry. 2014;19(10):1060–1070. doi:10.1038/mp.2014.91

86. Powell SB, Weber M, Geyer MA. Genetic Models of Sensorimotor Gating: Relevance to Neuropsychiatric Disorders. Curr Top Behav Neurosci. 2012;12:251. doi:10.1007/7854_2011_195

87. McGee a W, Topinka JR, Hashimoto K, et al. PSD-93 knock-out mice reveal that neuronal MAGUKs are not required for development or function of parallel fiber synapses in cerebellum. J Neurosci. 2001;21(9):3085–3091. doi:21/9/3085 [pii]

88. AR T, S M. Midbrain circuits of novelty processing. Neurobiol Learn Mem. 2020;176. doi:10.1016/J.NLM.2020.107323

