## Supplementary material for "Selective behavioural impairments in mice heterozygous for the cross disorder psychiatric risk gene *DLG2*": 051021 Pass Supplemental Information

**Supplementary Methods**

**Behavioural Analysis**

| Cohort | Numbers | Tasks |
|---|---|---|
| 1 | 46 mice (27 WT, 19 HET) | 2 day rotarod motor learning and performance, Acoustic startle and pre-pulse inhibition (PPI), protocol, and elevated plus maze (EPM) and open field anxiety tests. |
| 2 | 35 mice (18 WT, 17 HET) | 3 day rotarod motor learning protocol, and locomotor activity in activity box. |
| 3 | 24 mice (13 WT, 11 HET);<br>Control (4 WT, 3 HET) and rotarod-trained (8 WT, 8 HET) | cFos expression analysis after rotarod learning. |

**Supplementary Table 1. Experimental protocol used in the three cohorts of mice used during the study**. Two WT mice from Cohort 1 were euthanised during the course of experimentation due to illness, and one *Dlg2*$^{+/-}$ mouse (HET) was a consistent outlier (+/- 2SD from mean) and removed from statistical analysis. Tasks were conducted in order described.

**Anxiety Test: Elevated Plus Maze (EPM)**

The EPM was constructed from white Perspex in a cross formation arrangement of four arms: the two diametrically opposing 'open' arms with no walls and two 'closed' arms, 175 mm (L) x 78 mm (W), with 50 mm high walls. The maze was elevated 300 mm off the floor and illuminated at 15 lux. A computer running Ethovision Observer XT software (Noldus Information Technologies 3.0.15, Netherlands) was connected to a camera mounted above the maze tracking movement of the mouse during the trial. The Ethovision software tracked the mouse's position in the EPM and calculated analysis of time spent in predefined 'zones' within the maze, set up prior to testing by the experimenter and maintained across all trials.

Mice were placed in the nearest closed arm to the experimenter and allowed to explore freely for 5 minutes. The following behaviours were manually scored by the experimenter: number of head dips (downward movement of the rodent's head over the edge of an open arm), grooming and number of stretch attend postures (defined as an animal stretching forwards into the open arm whilst keeping its hindquarters in a closed arm). Time spent in the closed arms is indicative of anxious behaviour. Time spent in each maze zone, distance (m), velocity (m/s), latency to first entry into an open arm (s), and ethological parameters (grooming, head dips, stretch attend postures) were assessed using *t* tests.

**Anxiety Test: Open Field**

The open field apparatus consisted of a black Perspex floor (750 mm x 750 mm) with white Perspex walls (800 mm high), which was dimly illuminated (15 lux). A camera above the arena was connected to a computer running Ethovision Observer XT (Noldus Information Technologies 3.0.15, Netherlands) software recorded the animal's position (17 frames/s). The software subdivided the arena into a central zone (400 x 400 mm in arena centre) and an outer zone, within 350 mm of the walls. Mice were consistently placed into the closest corner of the arena to the experimenter, consistently facing the same wall. Animals could freely explore for the duration of the session (10 minutes). The main measures calculated were the duration of time spent in the central zone (the most exposed and therefore the most aversive part of the apparatus) and outer zone. The velocity (m/s) and total distance travelled (m) were recorded as an indices of activity. All measures were assessed by *t* tests.

**Genotyping Protocol**

Ear punches were taken during initial animal identification post weaning. Further tail tip biopsies were taken post-mortem for confirmation. All samples stored at -20°C. The Qiagen DNeasy Blood and Tissue Kit (Qiagen, Manchester, UK) was used as per the standard manufacturer's instructions to extract genomic DNA. Either 1 single ear-punch or approximately 0.6cm of tail tissue was lysed overnight at 56$^{o}$C in 180 µl ATL buffer and 20 µl proteinase K. 200 µl of AL buffer and 96-100% ethanol were added then all liquid was transferred to a spin column and centrifuged at 8000 rpm for 1 minute. The membrane was washed with AW1 and AW2 buffers and then the column was centrifuged at 14,000 rpm for 3 minutes to dry the membrane. 200 µl AE buffer was added directly to the membrane and incubated at room temperature for 1 minute. A final centrifugation step at 8000 rpm for 1 minute eluted the DNA, which was then stored at -20$^{o}$C. The Wellcome Trust Sanger Institute provided primer sequences (below) and a PCR cycling protocol for genotyping. Master mixes totalled 20 µl per reaction using the MyTaq™ DNA polymerase kit (Bioline, London, UK): 7.6 µl $ddH_20$, 4 µl 5x buffer, 0.8 µl reverse primer 1 and 2, 1.6 µl forward primer, 0.2 µl MyTaq™ and 5 µl DNA template.

Samples were run on a Biorad Thermal Cycler (T100 BioRad™, Herts, UK). Conditions for WT and mutant reactions were 94°C for 5 minutes, followed by 34 cycles of 94°C for 30 seconds, 58°C for 30 seconds and 72°C for 45 seconds, with a final extension for 5 minutes at 72°C and held indefinitely at 4°C.

**Supplementary Table 2**. PCR primer sequences and reaction temperature for genotyping of Dlg2t$^{m1a(EUCOMM)Wtsi}$ provided by the Wellcome Trust Sanger Institute.

| Primer | Reaction | Sequence (5' > 3') | Reaction Temp | Expected band size (bp) |
|---|---|---|---|---|
| Dlg2_42053_F | Wild type | CCAGAATGTACTTCAGCACCA | 58 | 312 |
| Dlg2_42053_R | | TGTGTGTATGTGTGGCTGTTT | | |
| Dlg2_42053_F | Mutant | CCAGAATGTACTTCAGCACCA | | 222 |

A 2% agarose gel was made with 1% Tris-acetate-EDTA (TAE) buffer (w/v) and SYBR Safe Gel DNA Stain (1:1000, ThermoFisherScientific, UK). Analysis was conducted by gel electrophoresis at 95 V for 45 minutes, with 10 µl PCR product loaded per well. Gels were visualised using an Omega Lum™ G imaging system (Apglegen, San Francisco, USA). WT animals were identified by one band (312 bp) whilst heterozygotes animals were identified by two bands (312 bp and 222 bp). Homozygous animals were identified by a mutant (222 bp) but not WT (312bp) bands.

**RT-qPCR: cDNA Thermocycler Settings and qPCR Primer Sequences**

**Supplementary Table 3**. Thermal cycler program for cDNA synthesis

| Temperature | Time |
|---|---|
| 42°C | 75 mins |
| 80°C | 15 minutes |
| 8°C | ∞ |

### Supplementary Data

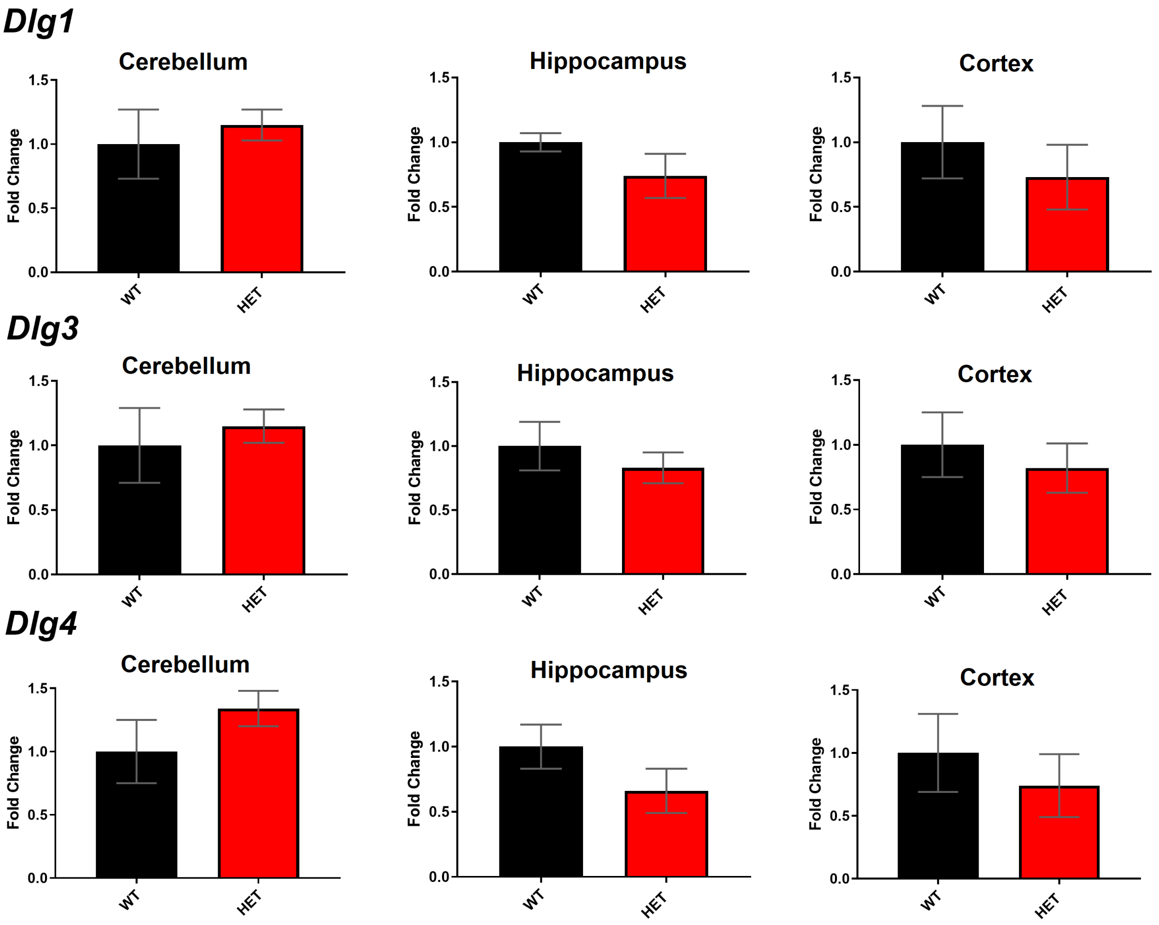


**Supplementary Figure 1. No difference in the regional expression of Dlg1, Dlg3 and Dlg4 in Dlg2$^{+/-}$ mice (HET) compared to WT as measured by RT-qPCR.** Error bars represent SEM. n = Cerebellum; 9 (WT), 10 (HET), hippocampus; 8 (WT), 9 (HET), and cortex (prefrontal cortex): 12 (WT), 12 (HET).

**Supplementary Table 4**. Statistical report for regional Dlg expression measured using RT-qPCR after t-test.

| **Gene/Region** | **Cerebellum** | **Hippocampus** | **Cortex** |
|---|---|---|---|
| *Dlg1* | $t_{(10.266)}$ 0.642, $p$ = 0.± 535 | $t_{(7.851)}$ -1.557, $p$ = 0.159 | $U$ = 50.00, $p$ = 0.219 |
| *Dlg3* | $t_{(12.645)}$ 0.936, $p$ = 0.367 | $t_{(15)}$ -1.107, $p$ = 0.286 | $U$ = 45.00, $p$ = 0.128 |
| *Dlg4* | $U$ = 43.00, $p$ = 0.631 | $t_{(15)}$ -1.755, $p$ = 0.100 | $U$ = 48.00, $p$ = 0.178 |

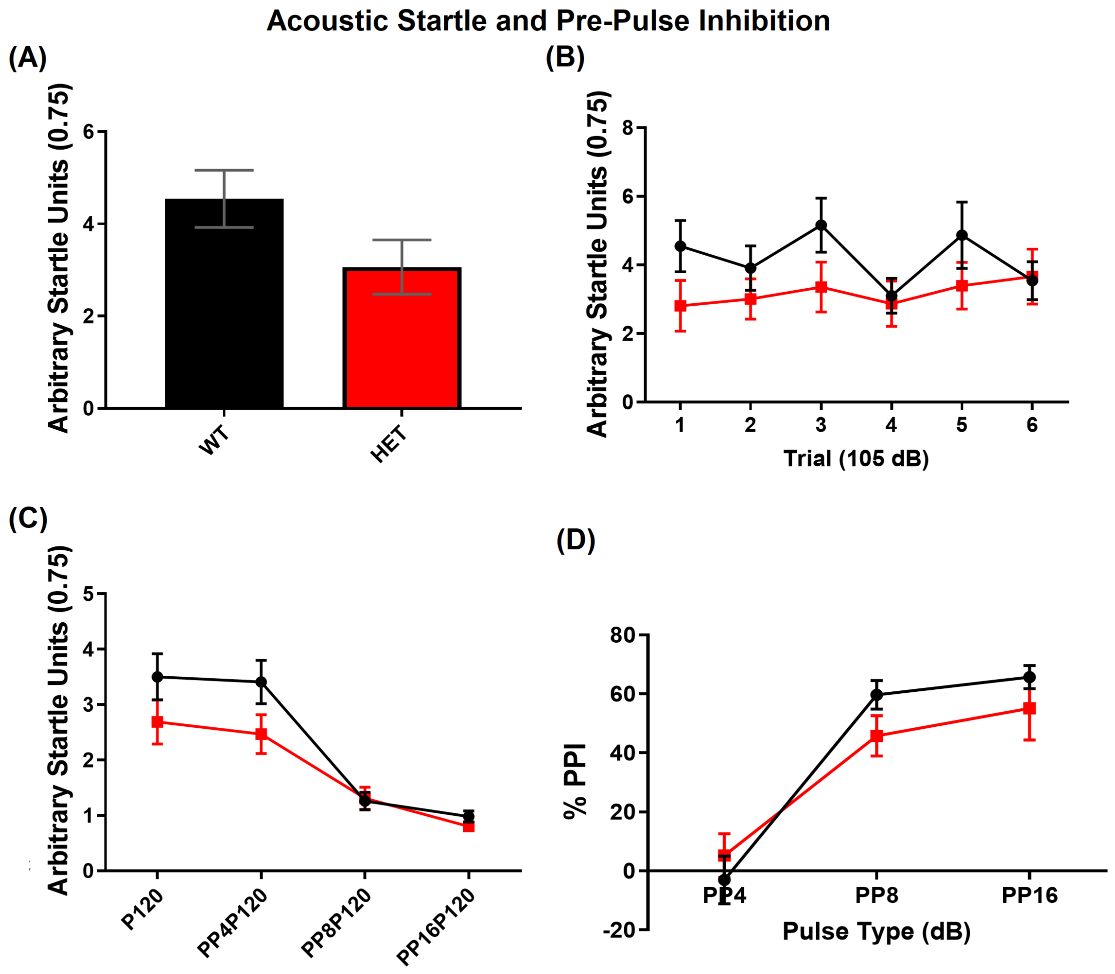


**Supplementary Figure 2. intact PPI and startle responses to a 105dB acoustic startle in *Dlg2*$^{+/-}$ mice.** (a) *Dlg2*$^{+/-}$ mice (HET) showed a trend towards reduced emotional reactivity compared to WT as measured as the averaged startle response to the first three startle stimuli at 105 dB ($t_{(41)} = 1.694$, $p = 0.098$, *t*-test). (b) However, no differences in startle responses between the genotypes were measured to the first 6 pulses at 105 dB (TRIAL: $F_{(5, 205)} = 1.975$, $p = 0.084$, GENOTYPE: $F_{(1, 41)} = 1.471$, $p = 0.232$, GENOTYPE x TRIAL: $F_{(5, 205)} = 1.485$, $p = 0.196$, mixed ANOVA). (c) There was a similar reduction of the startle response with an increasing intensity pre-pulse stimulus 70 ms before the 105 dB startle stimulus in both WT and HET mice. (PULSE: $F_{(1.660, 68.050)} = 49.730$, $p < 0.001$, GENOTYPE: $F_{(1, 41)} = 2.143$, $p = 0.151$, GENOTYPE x TRIAL: $F_{(1.660, 68.050)} = 2.219$, $p = 0.125$, mixed ANOVA). (d) Both HET and WT mice exhibited similar increases in startle response inhibition (% PPI) with pre-pulse intensity when expressed as a percentage of the 105 dB pulse-alone (P105) (PULSE: $F_{(1.643, 67.370)} = 55.852$, $p < 0.001$, GENOTYPE: $F_{(1, 41)} = 0.569$, $p = 0.455$, PULSE x GENOTYPE: $F_{(1.643, 67.370)} = 1.918$, $p = 0.162$, mixed ANOVA). PP4P105, PP8P105, and PP16P105 = pre-pulse 4dB, 8dB and 16 dB above background noise (70dB), respectively. Error bars are SEM. n = 24 (WT), 19 (HET).

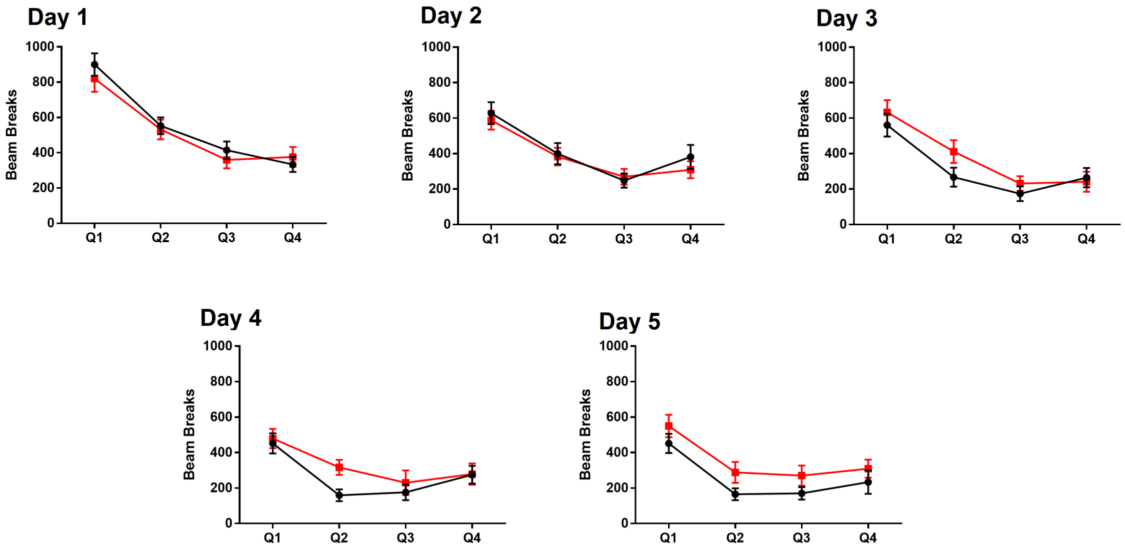


**Supplementary Figure 3. Within-session habituation to a novel context is normal in *Dlg2*$^{+/-}$ mice.** (a) Both WT and *Dlg2*$^{+/-}$ mice (HET) mice showed similar decreases in beam breaks per 30 min quantile (Q1-Q4) measuring locomotor activity in a 2-hour exposure to a locomotor activity box for 5 consecutive days (Days 1-5). There was an effect of Day ($F_{(4, 132)} = 31.266$, $p < 0.001$, mixed ANOVA) and Quantile ($F_{(2.006, 66.194)} = 83.879$ $p < 0.001$, mixed ANOVA), but not genotype ($F_{(1, 33)} = 0.229$, $p = 0.636$, mixed ANOVA), nor any two or three way interactions (*data not shown*). Analysing each day independently there was a reduction of activity (Day1, $F_{(2.297, 78.096)} = 99.747$, $p < 0.001$; Day2, $F_{(3,102)} = 44.161$, $p < 0.001$; Day 3, $F_{(3, 102)} = 41.235$, $p < 0.001$, Day 4, $F_{(3, 102)} = 16.684$, $p < 0.001$; Day 5, $F_{(3, 102)} = 23.397$, $p < 0.001$, mixed ANOVA) indicating intact within-session habituation. Data represent the mean ± SEM error bars. n = 18 (WT) 17 (HET).

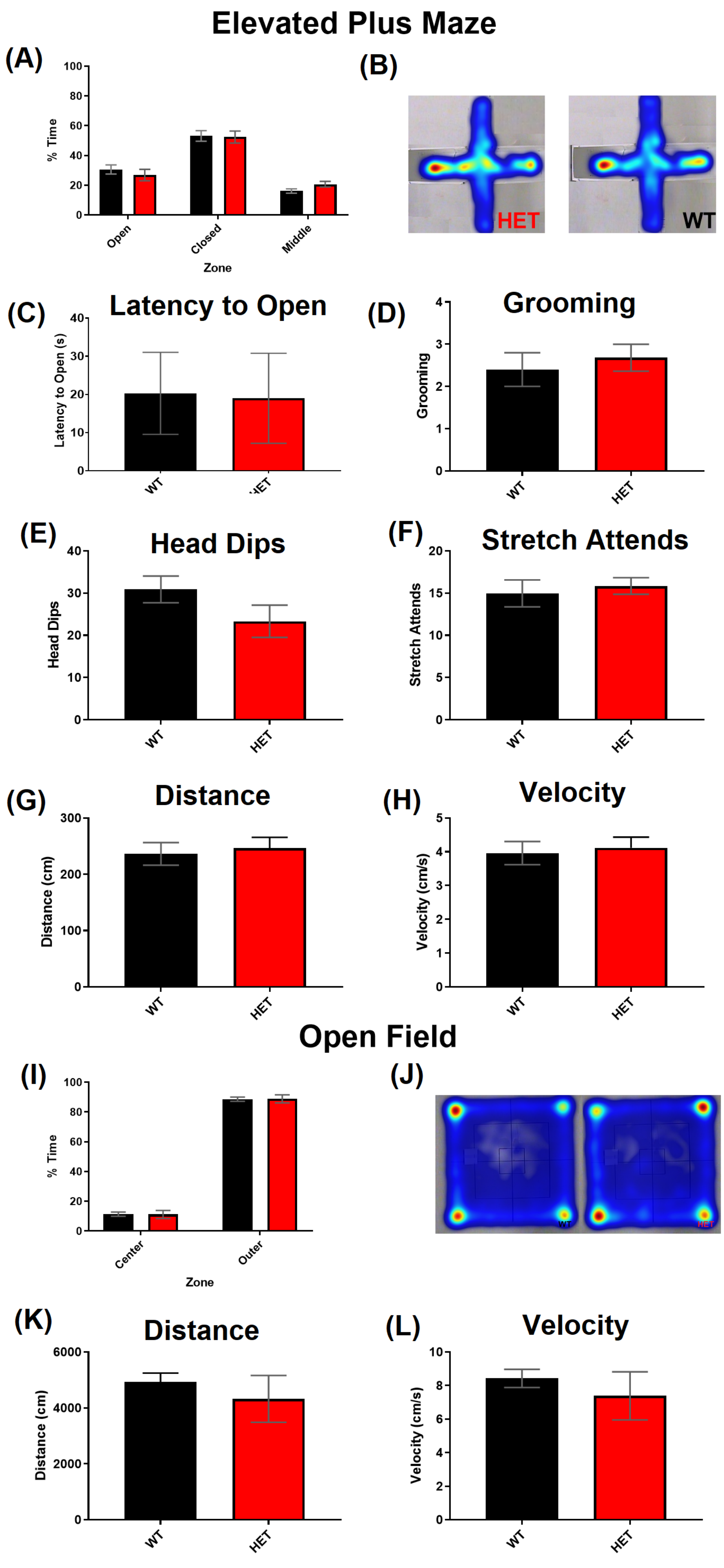

Elevated Plus Maze
(A)
% Time
100
80
60
40
20
0
Open
Closed
Middle
Zone
(B)
HET
WT
(C)
Latency to Open
Latency to Open (s)
40
30
20
10
0
WT
(D)
Grooming
Grooming
4
3
2
1
0
WT
HET
(E)
Head Dips
Head Dips
40
30
20
10
0
WT
HET
(F)
Stretch Attends
Stretch Attends
20
15
10
5
0
WT
HET
(G)
Distance
Distance (cm)
300
200
100
0
WT
HET
(H)
Velocity
Velocity (cm/s)
5
4
3
2
1
0
WT
HET
Open Field
(I)
% Time
100
80
60
40
20
0
Center
Outer
Zone
(J)
(K)
Distance
Distance (cm)
6000
4000
2000
0
WT
HET
(L)
Velocity
Velocity (cm/s)
10
8
6
4
2
0
WT
HET

**Supplementary Figure 4: No difference in anxiety-associated behaviours in the *Dlg2*$^{+/-}$ mice (HET) compared to WT measured in either the elevated plus maze (EPM) (a-h) or open field test (i-l).** (a) The % total time spent in each zone of the EPM during the 5 minute trial. (b) Merged heat maps for each genotype. Warmer colours indicate greater time spent in the maze zone. (c) Latency of mice to first enter an open arm of the elevated plus maze. (d) Number of periods of grooming during the 5 minute EPM trial (e) Numbers of head dips. (f) Numbers of stretch attends. (g) Total distance travelled during the 5 minute trial. (h) Mean velocity of movement during the 5 minute trial. There was no difference in any measure during the open field task (i-l). Both genotypes exhibited thigmotaxis, spending most time in the outer zone and corners of the open field. (i) The percentage time spent in each zone of the arena during the 10 minute trial. (j) Merged heat maps for each genotype. Warmer colours indicate greater time spent in the maze zone. (k) Total distance travelled during the 10 minute trial. (l) Velocity of movement during the 10 minute trial. Data represent the mean ± SEM error bars. n = EPM: 25 (WT), 19 (HET), open field test: 25 (WT), 18 (HET). For open field 1 HET was a consistent outlier across the measures and was removed.

**Supplementary Table 5.** Statistical report for separate measures of behaviour in the EPM and open field anxiety tests using *t*-test.

| Task | Measure | Statistic |
|---|---|---|
| Elevated plus maze | % Time in open arms | $t_{(42)}$, = 0.780, $p$ = 0.440 |
| | % Time in closed arms | $t_{(42)}$ = 0.146, $p$ = 0.885 |
| | % Time in center | $U$ = 156.50, $p$ = 0.055 |
| | Latency to Open Arm (s) | $U$ = 191.50, $p$ = 0.276 |
| | Distance travelled (cm) | $t_{(36.4)}$ = -0.471, $p$ = 0.640 |
| | Velocity (cm/s) | $t_{(36.32)}$ = -0.445, $p$ = 0.659 |
| | Stretch attends | $U$ = 191.50, $p$ = 0.275 |
| | Grooming | $U$ = 185.50, $p$ = 0.204 |
| | Head dips | $U$ = 193.50, $p$ = 0.297 |
| Open Field | % Time in inner zone | $t_{(40)}$ -0.388, $p$ = 0.351 |

| | | |
|---|---|---|
| | % Time in outer zone | $t$ $(_{40})$ 0.388, $p = 0.351$ |
| | Distance travelled (cm) | $t$ $(_{40})$ 1.167, $p = 0.243$ |
| | Velocity (cm/s) | $t$ $(_{40})$ 1.167, $p = 0.241$ |
